# Light-harvesting in mesophotic corals is powered by a spatially efficient photosymbiotic system between coral host and microalgae

**DOI:** 10.1101/2020.12.04.411496

**Authors:** Netanel Kramer, Raz Tamir, Or Ben-Zvi, Steven L. Jacques, Yossi Loya, Daniel Wangpraseurt

**Author notes:** Corresponding author: Netanel Kramer < >.

## Abstract

The coral-algal photosymbiosis fuels global coral-reef primary productivity, extending from sea level to as deep as 150 m (i.e., mesophotic). Currently, it is largely unknown how such mesophotic reefs thrive despite extremely limited light conditions. Here, we show that corals exhibit a plastic response to mesophotic conditions that involves a spatially optimized regulation of the bio-optical properties by coral host and symbiont. In contrast to shallow corals, mesophotic corals absorbed up to three-fold more light, resulting in excellent photosynthetic response under light conditions of only ~3% of the incident surface irradiance. The enhanced light harvesting capacity of mesophotic corals is regulated by average refractive index fluctuations in the coral skeleton that give rise to optical scattering and facilitate light transport and absorption by densely pigmented host tissue. The results of this study provide fundamental insight into the energy efficiency and light-harvesting mechanisms underlying the productivity of mesophotic coral reef ecosystems, yet also raise concerns regarding their ability to withstand prolonged environmental disturbances.

## Introduction

Scleractinian corals are the primary building blocks of coral reef ecosystems. Coral calcification involves the secretion of aragonite skeleton that provides the basis of the three-dimensional topography and complexity of coral reefs [1]. The underlying success of corals as ecosystem engineers is mainly due to the complex interaction that takes place between the coral hosts and their endosymbiont microalgae (family: Symbiodiniaceae). The coral-algal symbiosis is driven by solar energy and acts as the biological engine fueling the reef [2]. Corals have adapted to capture and maximize light under various environmental conditions, and it has been suggested that they are among the most efficient photosynthetic organisms at utilizing and converting light energy [3–6].

However, the symbiotic interaction is susceptible to anthropogenically mediated changes in environmental conditions [7]. Specifically, the coral-algal symbiosis is affected by periods of prolonged thermal stress, which can lead to a breakdown of the symbiosis and the visible paling of corals, known as coral bleaching [8]. Coral bleaching events have increased in frequency and duration over the last few decades, leading to an unprecedented decline of coral reefs worldwide [9]. The combined effects of elevated ocean temperatures, ocean acidification, and intensifying storms, have resulted in a decline in coral growth and functioning, due to reduced fecundity [10] and recruitment stock [11], reproductive synchronization breakdown [12], and coral disease outbreaks [13], thereby impairing the persistence of corals through environmental disturbances [14–16].

Most efforts to promote the recolonization of degrading shallow coral reefs have been focused on shallow-water corals [17]. More recently, it has been considered that mesophotic coral ecosystems (MCEs; > 30 m) are potential sources of replenishment and sinks for avoiding disturbances, since they could offer protection from the harmful environmental impacts encountered by their shallow-reef counterparts [18–20]. However, it has been shown that deep-water reefs can experience impacts from extreme storms and heatwaves, although not at the same frequency or intensity as in the shallow environments [21–23].

Coral morphological plasticity in response to environmental changes, is thought to enable corals to inhabit a wider range of environments and increase their ability to withstand disturbances [24–26]. Coral species that are distributed throughout a wide depth range (known as “depth-generalists”) adjust certain life-history traits in response to differences in irradiance [27]. It was found that light quantity affects coral morphology and growth rates [28, 29], community composition [30, 31], recruitment patterns [32–34], reproduction [35], and photobiology [27]. For instance, faster-growing species are found in well-lit shallow waters, while in deeper waters, decreasing levels of photosynthetically active radiation (PAR, 400-700 nm) typically result in reduced linear extension and coral calcification rates [28].

Although irradiance is fundamental for coral photosynthesis, excess light can easily result in photodamage to the photosynthetic apparatus [2]. Corals, therefore, employ a range of mechanisms to adjust light-harvesting and photosynthesis in response to the ambient light environment, most commonly by regulating their morphological structure [36,37], chlorophyll-*a* concentrations and/or cell densities [38,39]. Light-harvesting efficiency in corals is strongly controlled by the light scattering and absorption properties of coral host and symbionts. The light-scattering properties of coral host tissue and the aragonite skeleton play a key role in modulating the *in-hospite* light environment that controls photosynthesis [3,4]. While earlier coral optics studies have focused on the apparent optical properties (i.e. light field parameters), recent advances in experimental techniques enable the study of inherent optical properties (e.g., absorption and scattering coefficients) which depend on the material properties and structure of corals and are independent of illumination conditions [40–42]. For some corals, it has been shown that the high scattering of the living tissue traps light, while the low absorption and scattering of the skeleton redistribute the light that penetrates the living tissue, enabling the light to reach otherwise shaded living tissue [41].

The broad array of morphological forms found in corals indicates potentially important consequences in regard to regulation of the optical properties of the individual coral species [5,29,36]. In MCEs, corals can be found thriving at the PAR limits (0.1-1% of surface irradiance; see Tamir *et al*. 2019), which suggests that they have developed strategies to cope with and adjust to such extreme light habitats. Compared to shallow-water corals, those inhabiting mesophotic reefs exhibit unique characteristics that optimize photosynthetic efficiency [43–45]. Corals maximize their surface area at the morphological level to primarily laminar, plate-like morphologies allowing for a maximized light capture [43,46]. Furthermore, it has been suggested that fluorescent proteins can also promote the adaptation to low-light environments, by converting blue light into orange-red light, which can penetrate deeper within the coral tissues [43,47]. However, due to the previous inaccessibility of MCEs, our understanding of the bio-optical properties and ecophysiology of mesophotic corals are preliminary.

Here, we studied the ecophysiology and bio-optical properties of four widely depth-distributed coral species from shallow (5-10 m) and mesophotic (40-45 m) depths in the Gulf of Eilat/Aqaba, Red Sea. We aimed to elucidate the bio-optical mechanisms that enable corals to adapt to low-light environments, and hypothesized that corals in mesophotic environments would display bio-optical properties optimized to absorb low-light. Specifically, we employed a combination of techniques to study the photophysiology, *in-vivo* light field parameters, and the inherent optical properties of corals. The results of this study present an explanation of the processes that drive the photobiology and ecophysiology of mesophotic corals and shed light on how mesophotic corals may respond to environmental stress.

## Results

### Biometric assays

Algal pigmentation and density varied among species and within depths, with mesophotic specimens exhibiting an increase in pigmentation, as visible in the darker colored tissues (Fig. 1). Algal symbiont density (cells cm^-2^) was on average three-fold higher in the mesophotic corals than in their shallow counterparts (MEPA, *p* < 0.001; Fig. 2a), with the highest cell count measured in the mesophotic *P. lobata* at 6.57×10^5^ ± 1.11×10^5^ cells per cm^2^. Likewise, chlorophyll-*a* content (μg cm^-2^) was enhanced in mesophotic vs. shallow specimen (0.41-2.75 vs 0.014-0.94 μg cm^-2^, respectively; MEPA,*p* < 0.01; Fig. 2b). Overall, chlorophyll-*a* content per cell (pg cell^-1^) was higher in mesophotic corals (MEPA,*p* < 0.05; Fig. 2c), except for *S. pistillata* which exhibited similar concentrations between depths (*Hg* = −0.03 [C_I95%_ - 0.99; 0.99]).

**Figure 1.**
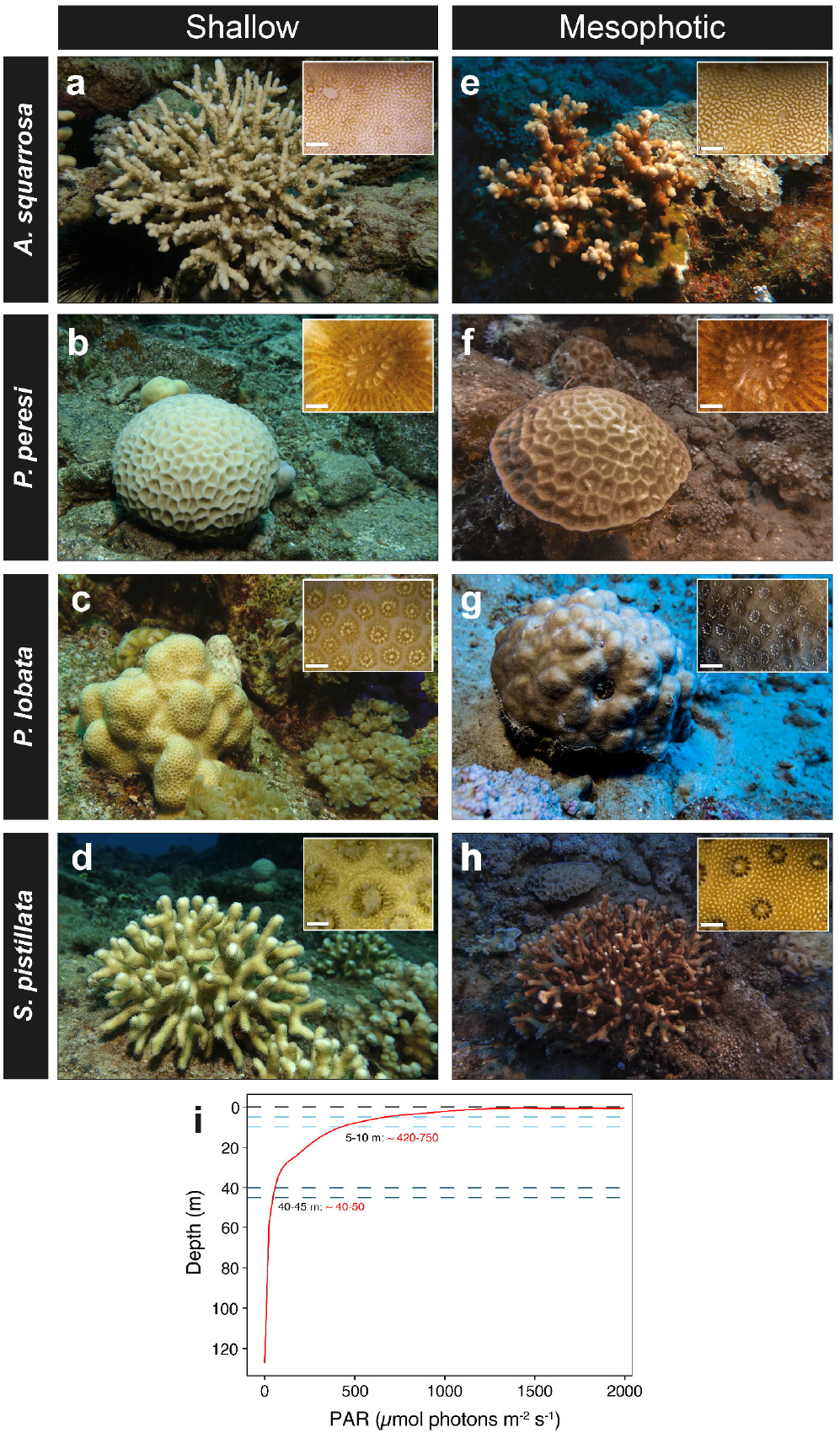
Morphotypes of depth-generalist corals and PAR profile. Studies species from shallow (5-10 m; **a-d**) and mesophotic (40-45 m; **e-h**) depths, at colony level and polyp-coenosarc level (insets; scale bars = 0.5 and 1 mm in branching and massive corals, respectively). Insets were taken using the same optical microscope settings. (**i**) Mid-day photosynthetic active radiation (PAR; μmol photon m^-2^ s^-1^) as a function of depth (m) in February, measured off the interuniversity institute for marine sciences in Eilat. Shallow (5-10 m; *light-blue*) and mesophotic (40-45 m; *dark-blue*) are shown as horizontal dotted lines, with a corresponding range of PAR values (*red*).

**Figure 2.**
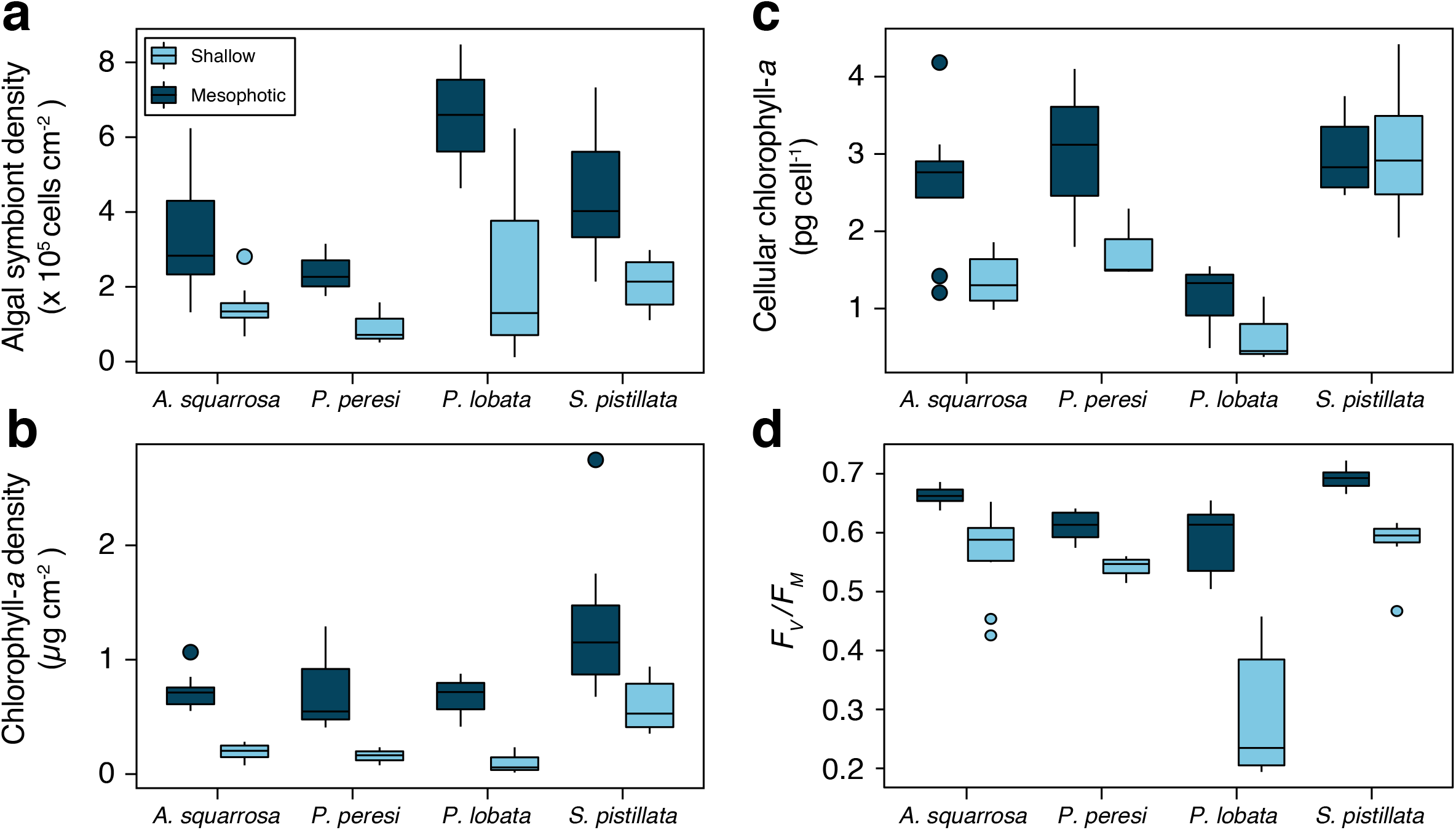
Biometric assays. Microalgal symbiont density (**a**), chlorophyll-*a* density (**b**), cellular chlorophyll-*a* (**c**), and *F*_v_/*F*_m_ (**d**) for four depth-generalist coral species from shallow water (5-10 m; *light-blue*) and mesophotic water depths (40-45 m; *dark-blue*). Box plots depict the median (horizontal line), interquartile range (first and third quartiles), and whiskers as ±1.5 interquartile range with dots representing outliers (*n* = 9-27 repetitions per species per depth).

### Photosynthetic parameters

The maximum quantum yield of PSII (*F*_v_/*F*_m_) was significantly higher for mesophotic species (MEPA, *p* < 0.05) compared to their shallow-water counterparts (*F*_v_/*F*_m_ = 0.50-0.72 and 0.19-0.65, respectively; p < 0.05), with shallow *P. lobata* exhibiting 50% lower *F*_v_*/F*_m_ values compared to all other shallow-water specimens (Fig. 2d). Areal net photosynthesis (μmol O_2_ cm^-2^ h^-1^) differed between branching species, as well as between depths within species (MEPA, *p* < 0.05). On average, the light-use efficiency (*α*) of areal net photosynthesis was three-fold higher for mesophotic corals compared to shallow-water corals (MEPA, *p* < 0.05; Fig. 3a, e). *S. pistillata* corals displayed the largest differences in *α* between depths, with values nine-fold higher in the mesophotic specimen than in their shallow-water counterparts (6.43×10^-2^ ± 3.33×10^-2^ vs 7.43×10^-3^ ± 1.66×10^-3^, respectively, Fig. 3a). Areal *P_MAX_* was enhanced for mesophotic corals compared to shallow ones, as can be seen for example in the two-fold higher *P_MAX_* in mesophotic *A. squarrosa* corals (*Hg* = 1.63 [CI_95%_ 0.774; 3.18]). Moreover, *P_MAX_* for mesophotic corals was achieved at almost one order of magnitude greater than in their ambient light environment (i.e., ~300 vs ~45 μmol photons m^-2^ s^-1^, Fig. 1i). All mesophotic specimens displayed significantly lower *EK* values than their shallower counterparts (MEPA, *p* < 0.01). In contrast to areal photosynthesis, cell-specific maximal rates in shallow *S. pistillata* exceeded those of their mesophotic counterparts by 30% (Table S1). The normalization for cell density also led to similar *P_MAX_* values between shallow and mesophotic *A. squarrosa* (*Hg* = 0.66 [CI_95%_ −0.66; 1.82]). The normalization of O_2_ production per unit cellular chlorophyll-*a* (μmol O_2_ pg chl-*a*^-1^ s^-1^) resulted in over two-fold greater *α* values for mesophotic corals (Table S1) and approximately 35% higher *P_MAX_* for shallow corals (MEPA, *p* < 0.05; Fig. 3c,g).

**Figure 3.**
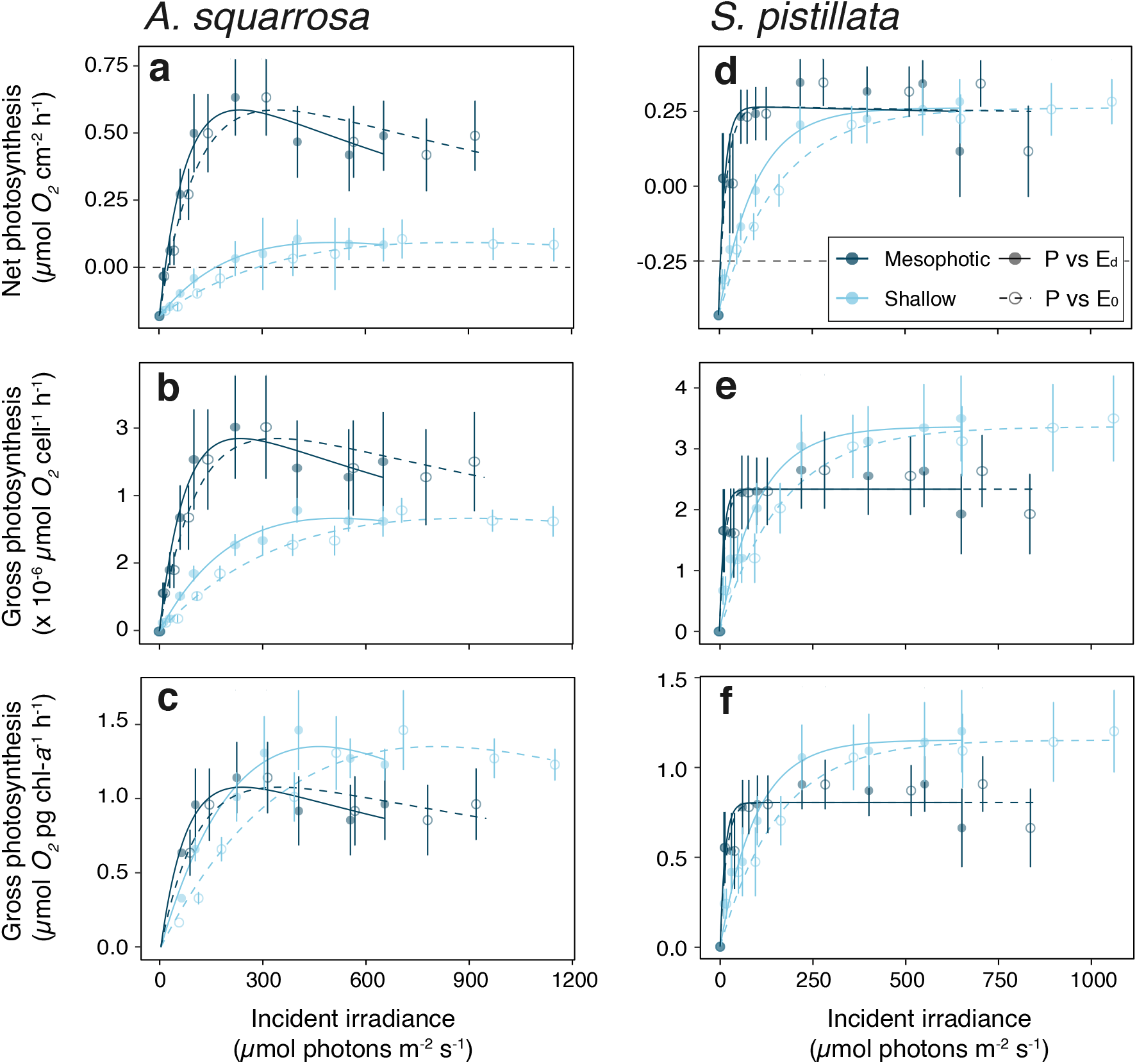
O_2_ production for *A. squarrosa* (a-c) and *S. pistillata* (e-g) from shallow (5-10 m; *light-blue*) and mesophotic (40-45 m; *dark-blue*) depths. O_2_ production is shown as areal net photosynthesis (μmol O_2_ cm^-2^ h^-1^; **a, e**), with gray dashed line represents 0 net photosynthesis; gross photosynthesis per algal cell (μmol O_2_ cell^-1^ h^-1^; **b, f**); and gross photosynthesis per chlorophyll content (μmol O_2_ pg chl-*a*^-1^ h^-1^; **c, g**). Each photosynthetic measurement (*P*) was performed as a function of the downwelling photon irradiance (*E_d_*; solid lines and filled circles) and of scalar photon irradiance (*E_0_*; dashed lines and hollow circles) spanning eight irradiance levels. Curves were fit using a double exponential decay model [61]. Data points represent mean ± standard error (*n* = 9 repetitions per species per depth). Note that the *x,y* scales for *A. squarrosa* and *S. pistillata* have been adjusted for clarity.

*P* vs *E_d_* was corrected for the *in vivo* photon scalar irradiance (*E_0_*; PAR integrated over 400-700 nm) for all the photosynthetic parameters. This correction resulted in an over 50% increase in *E_K_* values (*E_d_*; Table S1). For example, in shallow *A. squarrosa* corals, light saturation levels of *P* vs *E_0_* reached ~180% of the *P* vs *E_d_* (368.43 ± 79.40 and 206.73 ± 43.47 μmol photons m^-2^ s^-1^, respectively), whereas, in mesophotic corals, *E_K_* reached ~140% of the incident downwelling irradiance (91.85 ± 5.04 and 65.19 ± 3.03 μmol photons m^-2^ s^-1^, respectively; Fig. 3). There was no significant trend in dark respiration between all species and depths (Table S2; Fig. 3a, e). For massive coral species, no clear differences were found in photosynthetic parameters (Table S1).

### Bio-optical properties

Scalar irradiance (*E_0_*) at the coral tissue surface differed among species and depths for both tissue areas (i.e., coenosarc and polyp). Shallow corals exhibited 22% and 33% higher *E_0_* at 675 nm than mesophotic corals for coenosarc and polyp areas, respectively (MEPA, *p* < 0.001; Fig. 4a-d). Scalar irradiance was consistently higher over the coenosarc tissue than over the polyp tissue in both shallow and mesophotic *S. pistillata* corals (Fig. 4d), whereas *E_0_* differences between these areas in *P. lobata* varied only between depths, i.e., coenosarc was higher in shallow corals and lower in their mesophotic counterparts (Fig. 4c). Attenuation of PAR from tissue surface towards the skeleton was greater in mesophotic corals and was most pronounced in *P. peresi*, with *E_0_* reaching down to 24% of *E_d_* (Fig. S1). *E_0_* at the tissue-skeleton interface was approximately two-fold higher for shallow corals compared to mesophotic corals (MEPA, *p* < 0.001), ranging from 49.00 ± 0.88% to 158 ± 4.15% (mean ± SE; at 675 nm) of the incident downwelling irradiance. Intra-tissue measurements were not performed in *P. lobata* due to their extremely thin tissue.

**Figure 4.**
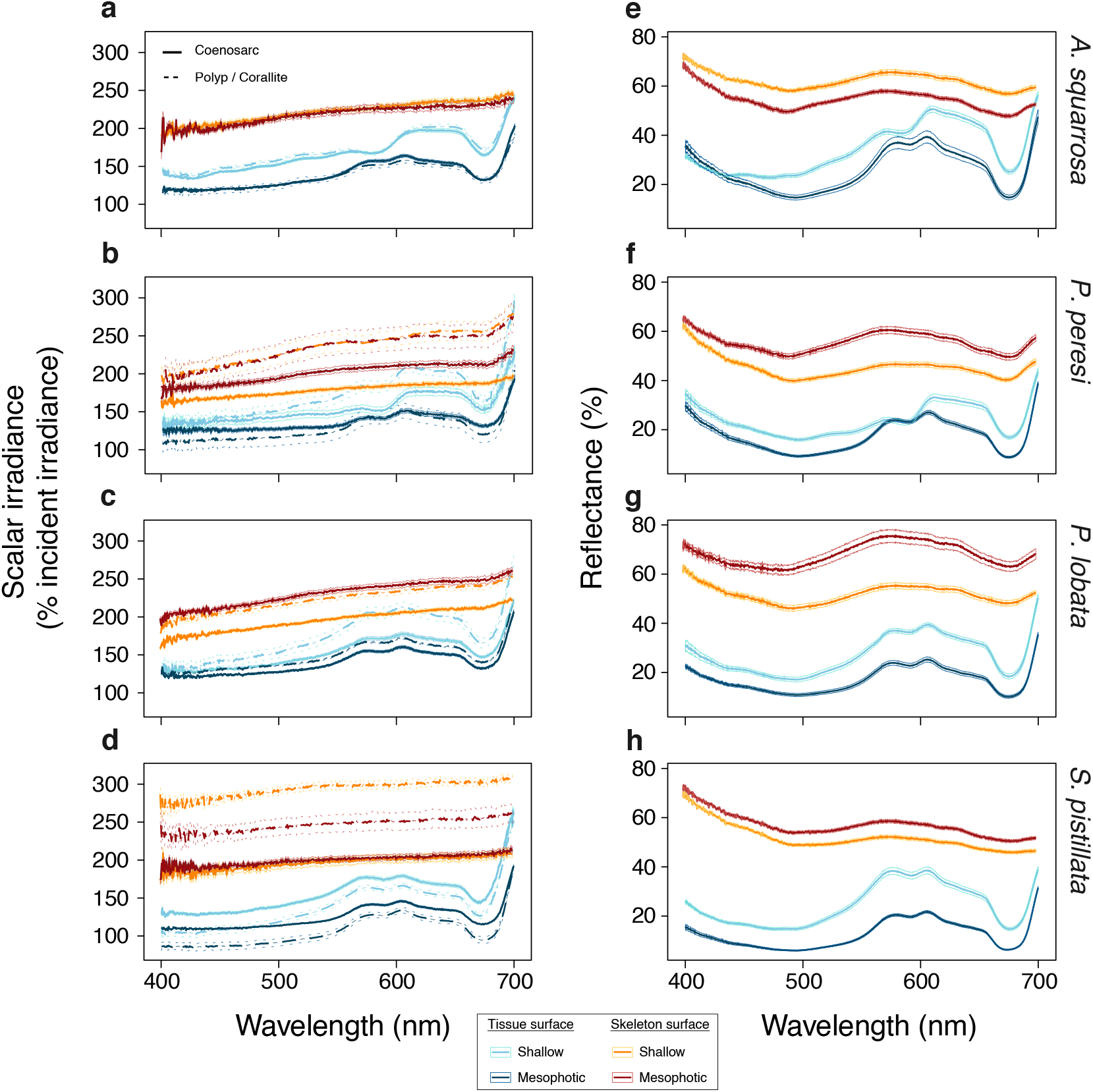
Apparent optical properties at 400-700 nm. Coral scalar photon irradiance (*E_0_*; % incident irradiance) at the tissue and skeleton surfaces, measured over the coenosarc (solid lines) and the polyp/corallite (dashed lines) areas (**a-d**). Normalized spectral reflectance of light (*Rd*(*λ*); %) over the coral tissue and skeleton surfaces (**e-h**). Tissue surfaces are colored in blue shades (*light-blue* and *dark-blue* for shallow and mesophotic corals, respectively), and skeleton surfaces are colored in warm shades (*orange* and *dark-red* for shallow and mesophotic corals, respectively). Data are mean (thick lines) ± standard error (thin lines); *n* = 15-45 repetitions per species per depth.

Skeleton scalar irradiance over corallite (polyp skeleton) areas of shallow and mesophotic corals were higher than over the coenosarc (MEPA, *p* < 0.001). For example, shallow *S. pistillata* corals exhibited nearly 50% higher skeleton scalar irradiance over corallite than over coenosarc, contrasting the pattern observed for intact corals (Fig. 4d). Compared to mesophotic corals, both the shallow massive coral species exhibited significantly 15% lower scalar irradiance over coenosarc (*P. peresi: Hg* = 1.95 [CI_95%_ 1.56; 2.34]; *P. lobata: Hg* = 1.81 [CI_95%_ 1.6; 2.04]), while no significant differences were found in the branching corals between depths.

The spectral diffuse reflectance (%) of corals varied among species, depths, and between live (tissue) and skeleton surfaces (MEPA, *p* < 0.001; Fig 4e-f). Tissue reflectance was systematically higher for shallow corals and ranged between 14.80 ± 0.26% to 25.20 ± 0.36% compared to 6.36 ± 0.10% to 14.70 ± 0.40% for their mesophotic counterparts (in λ = 675nm; Fig. 4e, f). Excluding *A. squarrosa*, coral skeletons from the mesophotic reflected up to 57% more light than their shallow counterparts (*Hg* = −1.71 [CI_95%_ −1.91; −1.52]; Fig. 4e).

The extracted algal absorption coefficient (*μ_a_algae_*; cm^-1^) for the four coral species ranged between 0.09 to 1.18 cm^-1^ at 675 nm (Fig. 5a-d, S2; Table 1). A comparison of shallow and mesophotic *μ_a_algae_* revealed that for most mesophotic specimens *μ_a_algae_* was at least two-fold greater. The largest difference was found for *A. squarrosa* with an approximate eight-fold enhancement in *μ_a_algae_* for mesophotic versus shallow corals. Skeletal optical parameters were extracted from the radial attenuation of the fluence rate at 720 nm (Fig. S3). Skeletal absorption was approximately two-fold higher in shallow versus mesophotic branching corals and was highest in the shallow *P. lobata* displaying 0.51 cm^-1^ (Fig. S4). The skeletal reduced scattering coefficient at 720 nm was relatively similar between shallow and mesophotic *S. pistillata* (15.12 and 13.5 cm^-1^, respectively). In contrast, shallow *A. squarrosa, P. peresi*, and *P. lobata* demonstrated over two-fold higher *μ_s_*’ compared to the mesophotic corals. Monte Carlo simulations used the extracted IOPs to calculate tissue absorption and showed that mesophotic *A. squarrosa* and *P. lobata* corals absorbed over 80% of the flux in the tissue compared to 25-30% in the respective shallow corals. In contrast, this depth-dependent difference in flux absorption was less pronounced for *S. pistillata* and absorption was only about 15% greater in mesophotic corals (Table 1).

**Figure 5.**
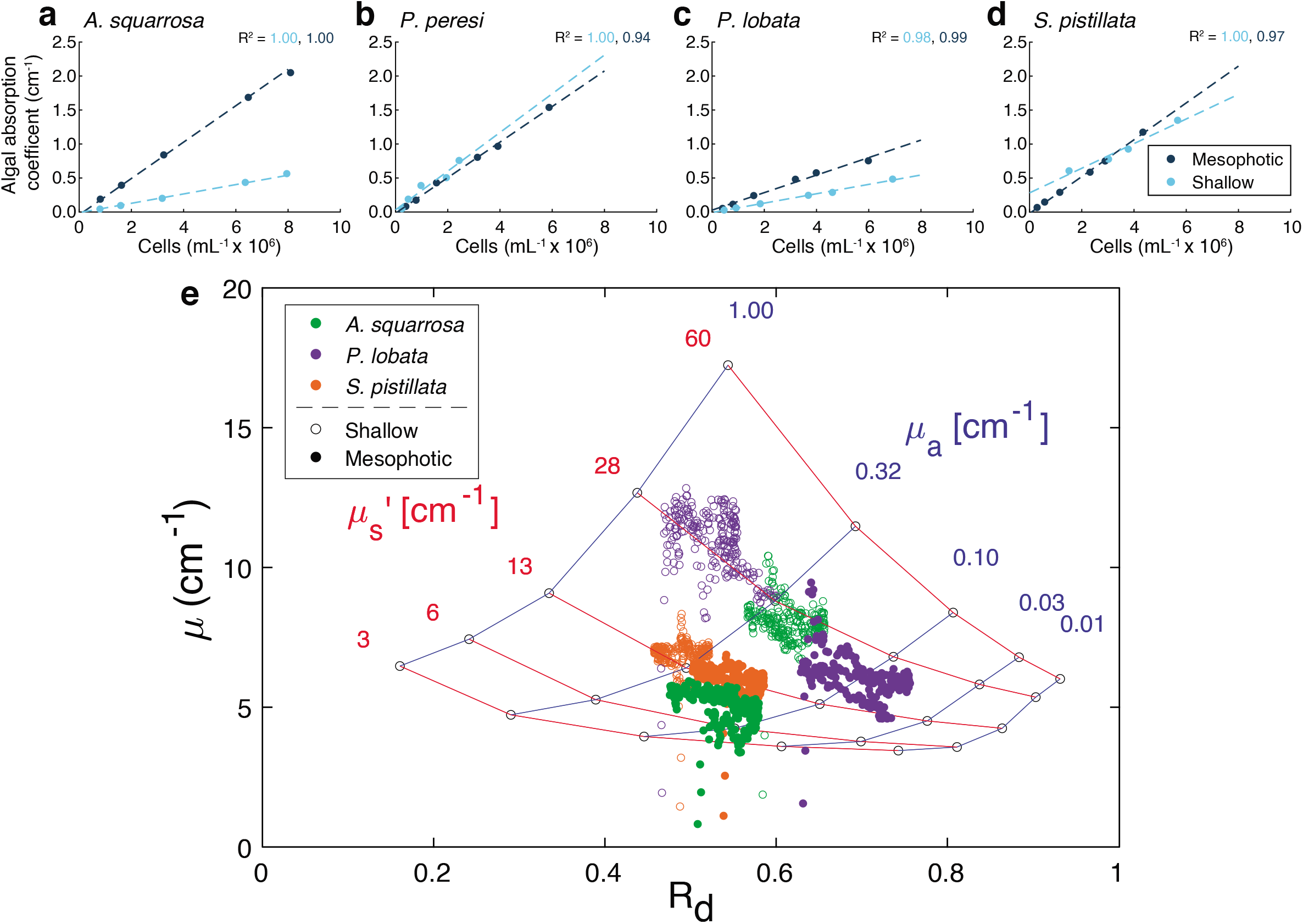
Inherent optical properties. Algal absorption coefficient *μ_a_algae_* (cm^-1^) of isolated microalgae at 675 nm as a function of microalgae density for shallow (*light-blue*) and mesophotic (*dark-blue*) samples (**a-d**). Coral skeletal scattering (*μ_s_’) and absorption coefficient* (*μ_a_*) between 500-750 nm (data points) plotted against measured *R_d_* for shallow and mesophotic (hollow and filled circles, respectively; colors denote species) (**e**).

**Table 1.**
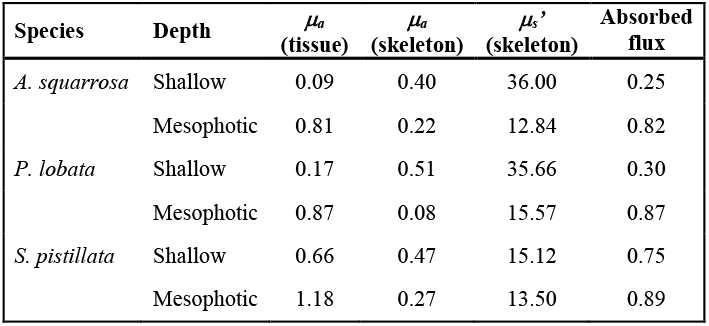
Inherent optical properties extracted from intact live corals (live at 675 nm) and bare skeletons (at 720 nm) using diffusion theory: tissue and skeletal absorption coefficient (*μ_a_*[cm^-1^]), and skeletal scattering coefficient (*μ_s_*’ [cm^-1^]). The predicted absorbed flux (watt per cm^-2^ per watt delivered) of the tissue affected by the optical parameters was derived from Monte Carlo simulations.

| Species | Depth | $\mu_a$<br>(tissue) | $\mu_a$<br>(skeleton) | $\mu_s'$<br>(skeleton) | Absorbed<br>flux |
| --- | --- | --- | --- | --- | --- |
| *A. squarrosa* | Shallow | 0.09 | 0.40 | 36.00 | 0.25 |
|  | Mesophotic | 0.81 | 0.22 | 12.84 | 0.82 |
| *P. lobata* | Shallow | 0.17 | 0.51 | 35.66 | 0.30 |
|  | Mesophotic | 0.87 | 0.08 | 15.57 | 0.87 |
| *S. pistillata* | Shallow | 0.66 | 0.47 | 15.12 | 0.75 |
|  | Mesophotic | 1.18 | 0.27 | 13.50 | 0.89 |

## Discussion

Understanding the ability of corals to grow and thrive under extreme low light is a key question in the study of mesophotic coral ecosystems [48]. In line with our hypothesis, the findings indicate that mesophotic corals have bio-optical properties optimized to absorb low-light. Our findings revealed up to three-fold enhanced light absorption by mesophotic corals (Fig. 5, Table 1) and an outstanding photosynthetic response under a range of light conditions (Fig. 3).

We found that for all mesophotic corals, except for *A. squarrosa*, PAR reflectance from coral skeletons was enhanced by up to 30% higher compared to shallow corals (Fig. 4). This shows that greater skeletal reflectance enhances light-harvesting by its photosymbionts [3], as exhibited by most mesophotic corals compared to their shallow counterparts (Fig. 4). We also detected a strong upregulation of light-absorbing pigments in all mesophotic coral tissues, due to both a higher algal cell density and enhanced chlorophyll-*a* per cell (Fig. 2), resulting in up to one order of magnitude higher algal absorption coefficients for mesophotic versus shallow-water corals (Fig. 5, Table 1). Such marked differences in algal absorption coefficient have not been previously characterized and indicate the presence of highly adapted light-harvesting complexes to low-light conditions. This is supported by earlier studies on mesophotic photobiology, demonstrating both an increase in the effective antenna size (i.e., antenna pigments) per photosynthetic unit (PSU) and an increase of PSUs per cell [44, 49].

Although the upregulation of photosynthetic pigments can naturally give rise to algal self-shading [3], microscopic light sensor measurements inside the tissue of mesophotic corals revealed that the light available at the tissue-skeleton interface is surprisingly high, leaving approximately one-fourth of the incident light after photon absorption (Fig. S1). It is possible, therefore, that the enhanced skeletal reflectance acts as an effective strategy counterbalancing the effects of self-shading, by upregulating the diffuse light available from the skeleton to compensate for higher cell densities.

We further found spatial differences in the microscale distribution of light between intact corals and bare skeletons (Fig. 4). For *S. pistillata* corals, tissue scalar irradiance was up to 20% lower for polyp than for coenosarc tissue, similar to previous measurements in other coral species characterized by small polyps [4]. The reduced light intensity for polyp tissues matches the spatial distribution of endosymbionts with higher cell densities in polyp tissues than in coenosarc tissues (see Fig. 1a,e). Interestingly, the irradiance distribution for the bare skeleton surface showed opposite patterns, indicating enhanced skeletal scattering within the polyp corallite than over the coenosarc. Therefore, enhanced light scattering within the corallite effectively assists in light dissipation to the dense microalgae inside the tissue. The high content of light-absorbing pigments within the polyp tissue (Fig. 4) suggests an increased biological activity of the coral polyp, and thus would require more light to support photosynthesis. Light availability within the corallite was strongly enhanced (Fig. 1S) which can be explained by intense scattering from the corallite skeletal walls toward its center [50], with additional light being transported to the polyp through the coenosteum [41]. Thus, the spatial distribution of the coral skeleton scattering and symbiont densities further support our hypothesis of a coral controlled finely tuned system of light scattering and light absorption.

Generally, both mesophotic *A. squarrosa* and *S. pistillata* corals utilized light more efficiently than their shallow counterparts, as indicated by steeper initial slopes of the *P-E* curves (Fig. 3a,d). For example, in *A. squarrosa* corals, the predicted net photosynthesis at a typical mid-day irradiance of 40-50 μmol photons m^-2^ s^-1^ in 45 m (Fig. 1i) was shown to be six-fold higher in mesophotic versus shallow depths (Fig. 3a). Still, we found species-specific differences in photosynthetic rates. Upon normalizing *P_n_* to algal cell density, *S. pistillata* revealed a typical light-shade response between depths (Fig. 3e), i.e., lower *P_MAX_* and *E_K_* values for mesophotic individuals [39,49]. In contrast, *A. squarrosa* did not follow this pattern and mesophotic corals exhibited higher *P_MAX_* and *E_K_* values when photosynthetic rates were normalized to cell density (Fig. 3b). However, when *P-E* curves were calculated based on chlorophyll-*a* content, *A. squarrosa* displayed a typical light-shade response, similar to *S. pistillata* (Fig. 3c,f).

The extraction of the inherent optical properties of corals allowed for a novel development of a probability distribution model that predicts the total flux absorbed within the photosynthetic coral tissue of shallow and mesophotic corals (Fig. 5, Table 1). The combination of increased reflectivity, high algal absorption coefficient, and low *μ_s_*’ (i.e., higher lateral spread of light) facilitated a strikingly higher total absorbed flux by mesophotic corals as compared with shallow ones. The mesophotic *A. squarrosa* and *P. lobata* showed approximately three-fold higher flux absorption than the shallower counterparts, whereas for *S. pistillata* there was only a ~20% difference between shallow and mesophotic specimen (Table 1). These findings indicate that in contrast to *A. squarrosa*, *S. pistillata* corals show light-regulated host modifications, as also supported by the pronounced differences in scalar irradiance between corallite and coenosarc areas, as well as between depths (Fig. 4d). Moreover, minor skeletal scattering differences between depths may be explained by micro-architectural modifications that compensate for changes in the skeletal material properties at mesophotic depths, as recently reported for this species [55]. This supports the notion that the ability of *S. pistillata* to adapt to different light regimes is not limited by its photosymbionts [53].

It is important to note that the energetic demands required to sustain coral growth at low-light can be further achieved through supplementation derived from heterotrophy (e.g., predation of zooplankton) [52]. However, since heterotrophic feeding is thought to be species-specific, the strategy of increased reliance on heterotrophy versus autotrophy with depth does not appear to be a primary trophic strategy for some depth-generalists, and particularly not in deep-water specialists [45]. The photosynthetic and growth efficiencies of the strictly mesophotic *Leptoseris* species, for example, were shown to be facilitated mainly by their skeletal optical geometry [45].

The advantage of low-light optimized bio-optical properties in mesophotic corals may prove to be their weakness in times of thermal stress. Although considered a relatively stable environment, MCEs can experience extreme heat-waves that lead to coral bleaching [22]. During bleaching in shallow corals, it has been shown that endosymbiotic algae are exposed to enhanced irradiance from skeletal backscattering, which can further stimulate symbiont loss, a hypothesis referred to as the “optical-feedback loop” [42,50]. Different factors have been shown to be related to enhanced bleaching susceptibility, such as lower *μ_s_*’ [42,54], enhanced skeletal reflectivity [50], and high photosymbiont densities [56], all of which have been demonstrated here by the mesophotic specimen (Fig. 3, 5, 6). Hence, the combination of these factors suggests that mesophotic corals are more susceptible to bleaching than their shallower counterparts. In an era of rapid climate change, it is therefore critical to assess the effects of thermal stress on mesophotic corals.

Thermal stress, however, is only one of many contemporary threats that coral reefs must face in a changing climate. Ocean acidification, in particular, was shown to impair the capacity of corals to build their skeletons through calcification [57,58]. An effective dissipation of light from the coral relies on a proper balance between the skeletal scattering and absorption, however, increased porosity and skeletal deformation caused by ocean acidification [59] are likely to disturb this balance. The light-harvesting process achieved by skeletal light scattering may become compromised as a result, leading to far-reaching ecological consequences caused by the coral’s inability to regulate light and impaired mechanical integrity. Given that corals are the main bioengineers in coral reefs, a reduction in coral growth will diminish the structurally complex habitat needed for numerous species [60]. Unfortunately, in times of change, this negative effect may be more pronounced in MCEs, since mesophotic corals could be more vulnerable and express limited and/or slower adaptability [22]. Notwithstanding the importance of corals as bioengineers, research focusing on the effects of altered environmental parameters on the bio-optical properties of corals has not received sufficient study to date, and more research is needed to determine the impact of projected near-future acidification levels, coupled with other significant environmental factors.

In conclusion, we present bio-optical mechanisms employed to sustain coral growth in extreme low-light environments, and expand the current knowledge on mesophotic photobiology. Light-harvesting in mesophotic corals is facilitated by the combination of pigment upregulation and enhanced skeletal reflectance. The light-harvesting strategies employed by mesophotic corals enhanced the absorbed flux by up to three-fold compared to their shallow counterparts, suggesting a spatially efficient photosymbiotic system. The results of this study further suggest that such light harvesting strategies make mesophotic corals specifically susceptible to environmental change and highlight the importance of integrating bio-optics and ecology to predict the future response of coral reef ecosystems to climate change.

## Author contributions

N.K., Y.L., and D.W. conceived and designed the study; N.K., R.T., and D.W. collected the corals; N.K., R.T., O.B.Z, and D.W. performed research; N.K., S.L.J, and D.W. analyzed the data; N.K. wrote the manuscript with contributions and final approval from all authors.

## Acknowledgments

We thank the Interuniversity Institute for Marine Sciences in Eilat (IUI) for making their facilities available, and their staff for excellent assistance. O. Levy, D. de Beer, and T. Treibitz are thanked for lending equipment. This work was supported by the Israel Science Foundation (ISF) Grant No. 1191/16 to Y.L., the European Union’s Horizon research and innovation program under grant agreement No. 730984, and by the ASSEMBLE-Plus consortium grant to D.W.

## STAR METHODS

### RESOURCE AVAILABILITY

#### Lead Contact

Requests for further information and data should be directed to and will be fulfilled by the lead contact, Netanel Kramer

#### Materials Availability

This study did not generate new unique materials.

#### Data and Code Availability

All datasets generated and/or analyzed during the study are available from the lead contact upon request.

### EXPERIMENTAL MODEL AND SUBJECT DETAILS

#### Coral collection and maintenance

Mature intact colonies of four depth-generalist coral species were chosen for a comparative study between shallow and mesophotic water specimens. We selected two branching coral species (*Stylophora pistillata* and *Acropora squarrosa*), and two massive coral species (*Paramontastrea peresi* and *Porites lobata*) based on their contrasting skeletal morphologies and their occurrence across a large depth gradient [29,30]. Coral colonies (*n* = 3 per species per depth) were collected using recreational and technical diving at shallow (5-10 m) and mesophotic (40-45 m) depths, respectively, from the Gulf of Eilat/Aqaba, the Red Sea. Conspecific colonies were collected at least five meters apart to avoid sampling clonemates. Colonies were maintained in outdoor open-circuit seawater tables at the Interuniversity Institute (IUI) in Eilat. Mesophotic colonies were kept in separate seawater tables under a blue light filter (Lagoon blue, Lee filters, UK) creating the spectral composition and intensity of irradiance at 45 m depth, while shallow corals were exposed to ambient sunlight. Downwelling irradiance was monitored to ensure that the light levels reflected the ambient conditions in which corals had been collected.

### METHOD DETAILS

#### Biometric measures

To determine microalgal cell density and chlorophyll*-a* content, coral tissue was removed using an airbrush at high pressure with 0.2 μm filtered seawater. Tissue-stripped fragments were bleached in a 6% sodium hypochlorite solution for 24 hours, then rinsed in freshwater for 10 minutes, and left to air dry. The bleached skeletons would be used later in the optical analyses. The microalgae fraction was separated from the host tissue using a motorized homogenizer and centrifugation (5000 rpm for 5 min). Following isolation, samples were immediately stored in a −80°C freezer for later analyses. Microalgal cell counts were determined using a hemocytometer on five replicate micrographs (scaled 0.1 mm^3^). Cell counts were normalized to the coral surface area to quantify areal algal density (cells cm^-2^). Chlorophyll-*a* was extracted from the remaining algae using 100% cold acetone for 15 h at 4°C and quantified spectrophotometrically [62]. Chlorophyll-*a* was normalized to surface area (μg cm^-2^) and algae cell (pg cell^-1^). The volume and surface area of the coral subsamples were determined with micro-computed tomography (μCT). The x-ray scans were conducted with a Nikon XT H 225ST μCT (Nikon Metrology Inc., USA) at a resolution of 50-μm voxels. Quantitative analysis was performed using Dragonfly software (v. 2020.1, Object Research Systems (ORS), Inc.).

#### Chlorophyll-*α* fluorometry

The maximum quantum yield of PSII (*F*_v_/*F*_m_) was measured with an imaging pulse-amplitude modulated (PAM) chlorophyll-*a* fluorometer (Maxi-PAM, Walz Gmbh, Germany). Coral samples were dark-acclimated for 20 minutes prior to each measurement (*n* = 3 per sample). The measured light intensity was adjusted to yield *F_0_* values in the region of interest that equal about 0.1 [63].

#### O_2_ turnover

Photosynthesis-irradiance (*P-E*) curves were performed for individual samples (*n* = 3-9). Each sample was incubated in a sealed 270 ml acrylic metabolic chamber. The chambers were placed in a temperature-controlled bath (RTE 210, Thermo Neslab), with constant water flow at 22°C (i.e., ambient seawater temperature), and a magnetic stirrer maintaining water movement inside the chamber. A full-spectrum metal halide lamp (400 W, 5000 K, 50 Hz, Golden-Light, Netanya, Israel) was used to incubate the corals at incident downwelling irradiance (*E_d_*) regimes spanning 0 to 800 μmol photons m^-2^ s^-1^. *E_d_* was measured using a light meter (LI-250A, Li-Cor, Inc. Lincoln, NE, USA) connected to a cosine-corrected quantum sensor. O_2_ evolution was monitored within each chamber using O_2_ optodes connected to an O_2_ meter (ProODO Optical Dissolved Oxygen meter, YSI Inc., OH, USA). Areal net photosynthesis (*Pn*) was calculated from the difference between final and initial O_2_ measurements (ΔO_2_) for each session after 20 minutes under each light intensity. The *P-E* data were fit to a double exponential decay function to characterize the photosynthetic efficiency (*a*), the maximal photosynthesis rate (*P_MAX_*), and the minimum saturation irradiance (*E_K_*) [61]. Furthermore, photosynthetic rates were normalized to symbiont cell density and chlorophyll-*a* content to examine the role of the coral host optics in affecting photosynthetic efficiency. Cell-specific gross photosynthetic rates (μmol O_2_ cell^-1^ s^-1^) were based on the assumption that light respiration was up to 1.5-fold higher than dark respiration [50].

##### Apparent optical properties (AOPs)

Spectral scalar irradiance *E_0_(λ)* and diffuse reflectance *R_d_(λ)* were measured for intact corals and the bare skeletons of each individual. Measurements were performed in a black acrylic flow chamber. Fiber-optic scalar irradiance microprobes with a tip diameter of 50-100 μm (Zenzor, Denmark) were used to measure the surface and intra-tissue light microenvironment as described previously [4]. The microsensors were connected to a spectrometer (AvaSpec-UL2048XL, Avantes, USA) and data were recorded with commercial software (Avasoft 8.0, Avantes, USA). *E_0_(λ)* was normalized to the incident downwelling irradiance *E_d_(λ)*, which was measured under an identical configuration as the experimental coral measurements [4].

Spectral reflectance *R_d_(λ)* was measured with a flat-cut fiber-optic reflectance probe (diameter = 0.2 cm, Ocean optics, USA) connected to a portable spectrometer (JAZ, Ocean optics, USA). For each measurement, the probe was positioned at 5 mm from the coral/skeleton surface and at a 45° angle relative to the surface [64]. Incident irradiance was provided by a tungsten halogen lamp (Schott ACE 1, Germany) equipped with a collimating lens. Measurements were taken on five randomly chosen areas per coral. Experimental measurements were normalized against a measurement performed on a 99% diffuse reflectance standard (Spectralon, Labsphere USA). Although skeletons were bleached, the reflectance spectrum in the peridinin-chlorophyll-protein complex and chlorophyll-*a* wavebands (490-500 and 675 nm, respectively) were slightly affected by pigment residuals (presumably from remaining endolithic algae). Nevertheless, this did not affect the interpretation of the results.

##### Inherent optical properties (IOPs)

The transfer of light in corals is described by the radiative transfer equation (RTE). However, the RTE is difficult to solve analytically. In most scattering dominating systems, the RTE can be simplified and expressed as a diffusion dominated process, where optical energy diffuses according to the diffusion equation [65]. Farrell et al. (1992) developed a steady-state diffusion equation for light transport in a semi-infinite planar geometry, where the diffuse reflectance *R* leaving the boundary at a given distance ρ from the source is:

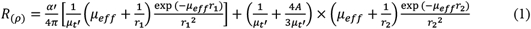

where *μ_eff_* is the effective attenuation coefficient:

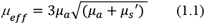

*a’* is the transport albedo:

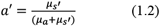

*μ_t_* is the total interaction coefficient:

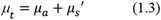

And *r_1_* and *r_2_* are radial distances of one mean free path inside the medium and above the medium where total fluence equals 0, respectively [66]. The optical properties *μ_s_’* and *μ_a_* uniquely determine the shape of the diffuse reflectance curve. By measuring the lateral spread (*r*) of reflected light (*R*) it is thus possible to predict unique values of *μ_s_*’ and the absorption coefficient *μ_a_* that generated the reflectance profile. The fitting procedure uses the fminsearch.m routine in Matlab (Mathworks, USA) which calls a multidimensional unconstrained non-linear minimization algorithm (Nelder–Mead) to minimize the sum of squares error [41,67]. Optical extraction of *μ_s_*’ and *μ_a_* were performed for intact corals at wavelengths of strong chlorophyll-*a* absorption (at 663 nm) as well as in the near-infrared (at 750nm), which is free from pigment absorption.

We extracted scattering (*μ_s_*’) and absorption (*μ_a_*) coefficients [68] from intact corals and skeletons by measuring the lateral spread of reflected light (*R_r_*). Measurements were performed with a flat cut light-emitting source fiber (diameter = 0.2 cm, Ocean optics, USA) connected to a tungsten halogen lamp (LS-1, Ocean Optics, USA) and a flat-cut light collecting fiber (diameter = 50 μm, Zenzor, Denmark) connected to a spectrometer (AvaSpec-UL2048XL, Avantes, USA). Both fibers were mounted on micromanipulators (Pyroscience GmBH, Germany and Märtzhäuser, Germany) and aligned parallel to each other at a minimum lateral distance of 4 mm perpendicular to and in direct contact with the coral surface. A stereomicroscope (SZ51, Olympus) was used to carefully position the fiber optic probes at the surface. *Rr* was measured at lateral steps of 1 mm to a maximum of 10 mm [67]. This procedure was repeated for five randomly chosen coenosarc areas for each coral sample. Measurements were conducted on the coenosarc, which had a more even topography and less contractile tissue compared with the polyp tissue, thus allowing repeated measurements and a more accurate estimate of horizontal light transfer. The lateral attenuation of light was matched to the predicted attenuation *pR_r_* based on diffusion theory [66] and starting values of *μ_s_’ μ_a_*. The coral surface architecture for *P. peresi* was very heterogeneous due to the deep corallite architecture, preventing reliable quantification of *R*(*r*), and the analysis was thus omitted.

Additionally, we characterized the algal cell-specific absorption coefficient (*μ_a_algae_*) of isolated symbionts independent of the host environment. To separate between the effect of algal absorption and that of algal scattering, diffuse reflectance measurements were performed in a strongly scattering dominated medium, such that any algal scattering can be regarded as negligible. Milk is a cost-effective strongly scattering-medium with known optical properties.

The lipid content of 100% whole milk is typically 4% lipids and the scattering of intralipid^™^ in 10% lipids is 100 cm^-1^ (at 600 nm). Therefore, the scattering of 100% whole milk is estimated to be:

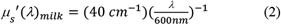

Reflectance measurements were performed with 100% whole milk and a 1:1 mixture of milk and algal serial dilutions (from 50% to 3%). Algal cell density was determined as described above in order to relate *μ_a_algae_* to cell density (mL^-1^). To extract *μ_a_algae_* [cm^-1^] diffusion theory and nonlinear least-squares fitting were used as described above to match predicted reflectance to experimentally measured reflectance.

#### Monte Carlo modeling of absorbed flux

To characterize differences in light absorption by coral tissues we developed a Monte Carlo simulation using the inherent optical properties determined via diffusion theory as input parameters (Fig. 6, Table 1). For each simulation the tissue was 1mm thick and had an absorption coefficient *μ_a_* that was determined by *μ_a_* algae at 675 nm. Tissue *μ_s_*’ was set to 5 cm^-1^ for all simulations. Details on the Monte Carlo approach can be found in Wangpraseurt et al. [41] and Jacques et al [67].

### QUANTIFICATION AND STATISTICAL ANALYSIS

All statistical analyses were performed using the R software (R Core Team 2020). To estimate ecophysiological and bio-optical variations in the studied species, we modeled the corresponding parameters (separate test for each parameter) as a function of species and depth, using mixed-effects permutational analysis (MEPA; 2000 permutations), with coral identity as a random effect. These analyses were run using the packages {nlme}[70] and {predictmeans}[71]. Pairwise comparisons were based on Hedge’s *g* (*Hg*) standardized effect size (preferred over Cohen’s *d* for small samples) with 95% confidence interval (CI) constructed from 5000 bootstrap samples, and significance was determined as CI not overlapping with zero (shallow depth as reference). This analysis was computed using the R package {dabestr}[72].

