## Supplementary for "Light-harvesting in mesophotic corals is powered by a spatially efficient photosymbiotic system between coral host and microalgae"

[1] *School of Zoology, Tel-Aviv University, Tel Aviv 69978, Israel*

### Supplementary Tables:

**Table S1.** Photosynthesis vs irradiance/energy (*P-E*) curve fitting parameters. Photosynthetic rates (*n* = 3-9 measurements per species per depth) are presented as:**1)** areal net photosynthesis **2)** gross photosynthesis per algal cell, and **3)** gross photosynthesis per chlorophyll content for the four depth-generalist coral species between shallow (5-10 m) and mesophotic (40-45 m) depths. The parameters were fitted and extracted from a double exponential decay model [1]: initial slope ($\alpha$); coefficient of determination range ($R_2$); maximum gross photosynthesis rate ($P_{MAX}$); and minimum saturation irradiance ($E_K$) calculated for the incident downwelling irradiance ($E_d$) and *in vivo* photon scalar irradiance ($E_0$). Standard errors are in parenthesis.

| | **Species** | **Depth** | **Slope ($\alpha$)** | **$R^2$** | **$P_{MAX}$** | **$E_K$ ($E_d$)** | **$E_K$ ($E_0$)** |
|---|---|---|---|---|---|---|---|
| **1. Areal net photosynthesis** | *A. squarrosa* | Shallow | $1.68\times10^{-3}$ ($5.16\times10^{-4}$) | 0.92 (0.04) | 0.30 (0.07) | 206.73 (43.47) | 368.43 (79.40) |
| | | Mesophotic | $1.19\times10^{-2}$ ($2.56\times10^{-3}$) | 0.92 (0.02) | 0.76 (0.15) | 65.19 (3.03) | 91.85 (5.04) |
| | *P. peresi* | Shallow | $8.40\times10^{-3}$ ($2.28\times10^{-3}$) | 0.89 (0.01) | 0.80 (0.08) | 99.13 (18.63) | 163.04 (41.32) |
| | | Mesophotic | $1.18\times10^{-2}$ ($1.16\times10^{-3}$) | 0.96 (0.00) | 0.89 (0.11) | 75.56 (1.63) | 105.76 (1.85) |
| | *P. lobata* | Shallow | $3.21\times10^{-2}$ ($2.21\times10^{-2}$) | 0.96 (0.01) | 2.33 (1.50) | 79.08 (8.59) | 127.02 (15.41) |
| | | Mesophotic | $5.56\times10^{-2}$ ($1.79\times10^{-2}$) | 0.96 (0.01) | 3.06 (1.43) | 51.33 (8.23) | 73.70 (10.29) |
| | *S. pistillata* | Shallow | $7.43\times10^{-3}$ ($1.66\times10^{-3}$) | 0.92 (0.02) | 0.63 (0.07) | 122.78 (30.86) | 198.73 (48.53) |
| | | Mesophotic | 6.43s$\times10^{-2}$ ($3.33\times10^{-2}$) | 0.86 (0.01) | 0.66 (0.02) | 35.55 (27.63) | 45.65 (35.44) |

| | Species | Depth | Slope ($\alpha$) | $R^2$ | $P_{MAX}$ | $E_K$ ($E_d$) | $E_K$ ($E_0$) |
|---|---|---|---|---|---|---|---|
| **2. Gross photosynthesis per algal cell** | *A. squarrosa* | Shallow | $1.00\times10^{-8}$ ($1.60\times10^{-9}$) | 0.91 (0.04) | $1.90\times10^{-6}$ ($1.92\times10^{-7}$) | 215.20 (41.30) | 380.52 (75.68) |
| | | Mesophotic | $4.52\times10^{-8}$ ($1.32\times10^{-8}$) | 0.89 (0.03) | $2.86\times10^{-6}$ ($7.73\times10^{-7}$) | 65.19 (3.03) | 97.19 (6.69) |
| | *P. peresi* | Shallow | $1.26\times10^{-7}$ ($8.74\times10^{-8}$) | 0.89 (0.01) | $1.09\times10^{-5}$ ($6.31\times10^{-6}$) | 99.13 (18.63) | 162.74 (41.63) |
| | | Mesophotic | $5.89\times10^{-8}$ ($1.70\times10^{-9}$) | 0.61 (0.00) | $4.55\times10^{-6}$ ($3.20\times10^{-8}$) | 75.56 (1.63) | 105.76 (1.85) |
| | *P. lobata* | Shallow | $3.44\times10^{-7}$ ($2.45\times10^{-7}$) | 0.93 (0.05) | $2.51\times10^{-5}$ ($1.71\times10^{-5}$) | 79.08 (8.59) | 127.02 (15.41) |
| | | Mesophotic | $9.16\times10^{-8}$ ($3.02\times10^{-8}$) | 0.77 (0.15) | $4.87\times10^{-6}$ ($2.14\times10^{-6}$) | 51.33 (8.28) | 73.70 (10.29) |
| | *S. pistillata* | Shallow | $4.38\times10^{-8}$ ($1.32\times10^{-8}$) | 0.92 (0.02) | $3.41\times10^{-6}$ ($6.14\times10^{-7}$) | 125.38 (31.05) | 202.84 (48.74) |
| | | Mesophotic | $2.78\times10^{-7}$ ($1.61\times10^{-7}$) | 0.86 (0.01) | $2.51\times10^{-6}$ ($4.39\times10^{-7}$) | 35.55 (27.63) | 45.65 (35.44) |
| **3. Gross photosynthesis per chlorophyll content** | *A. squarrosa* | Shallow | $8.16\times10^{-3}$ ($1.14\times10^{-3}$) | 0.92 (0.04) | 1.45 (0.14) | 208.18 (43.20) | 366.61 (79.67) |
| | | Mesophotic | $1.71\times10^{-2}$ ($4.28\times10^{-3}$) | 0.92 (0.02) | 1.08 (0.25) | 65.19 (3.03) | 91.85 (5.04) |
| | *P. peresi* | Shallow | $8.41\times10^{-2}$ ($5.79\times10^{-2}$) | 0.89 (0.01) | 7.26 (4.17) | 99.13 (18.63) | 162.10 (42.27) |
| | | Mesophotic | $2.56\times10^{-2}$ ($6.18\times10^{-3}$) | 0.96 (0.00) | 1.94 (0.51) | 75.56 (1.63) | 105.76 (1.85) |
| | *P. lobata* | Shallow | $4.06\times10^{-1}$ ($1.64\times10^{-1}$) | 0.96 (0.01) | 30.20 (10.76) | 79.08 (8.59) | 127.02 (15.41) |
| | | Mesophotic | $7.96\times10^{-2}$ ($1.11\times10^{-2}$) | 0.96 (0.01) | 4.25 (1.27) | 51.33 (8.28) | 73.70 (10.29) |
| | *S. pistillata* | Shallow | $1.59\times10^{-2}$ ($5.19\times10^{-3}$) | 0.92 (0.02) | 1.17 (0.20) | 122.78 (30.86) | 198.76 (48.54) |
| | | Mesophotic | $8.89\times10^{-2}$ ($4.55\times10^{-2}$) | 0.86 (0.01) | 0.87 (0.09) | 35.55 (27.63) | 45.65 (35.44) |

**Table S2.** Coral dark respiration rates (*R*, µmol $O_2$ cm$^{-2}$ h$^{-1}$). Data are means ± standard error in parenthesis (*n* = 3-9 measurements per species per depth).

| **Species** | **Depth** | ***R*** |
|---|---|---|
| *A. squarrosa* | Shallow | 0.12 (0.03) |
| | Mesophotic | 0.14 (0.03) |
| *P. peresi* | Shallow | 0.22 (0.03) |
| | Mesophotic | 0.09 (0.01) |
| *P. lobata* | Shallow | 1.32 (0.85) |
| | Mesophotic | 0.49 (0.11) |
| *S. pistillata* | Shallow | 0.25 (0.05) |
| | Mesophotic | 0.32 (0.11) |

**Supplementary Figures:**

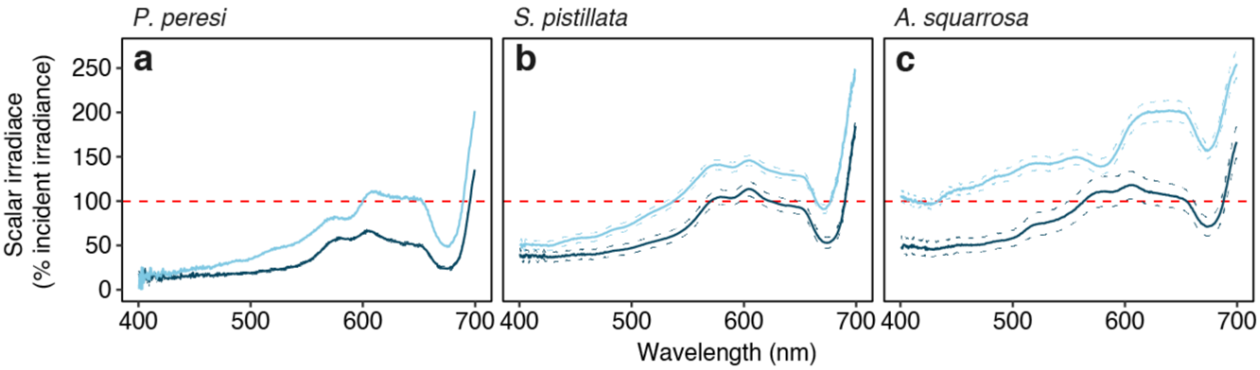


**Figure S1.** Spectral scalar irradiance close to the skeleton surface within the polyp tissue. Dotted red line represents 100% of incident irradiance ($E_d$). Spectral measurements of three species are colored by depth: shallow (5-10 m; *light-blue*) and mesophotic (40-45 m; *dark-blue*). Tissue thicknesses ranged from 400 µm to 900 µm. Data are mean (solid lines) ± standard error (dashed lines).

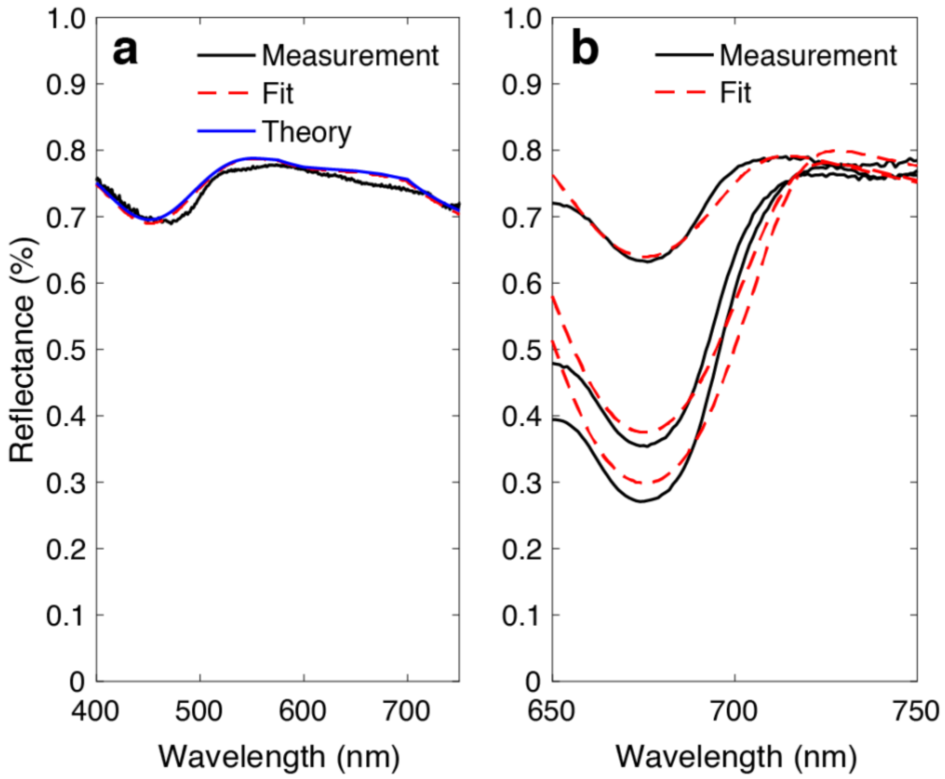


**Figure S2.** Principle of algal absorption coefficient extraction. Diffuse reflectance measurements from 100% whole milk (*black*), best fit predicted from diffusion theory (*red*), and theoretical prediction based on the scattering of intralipid (*blue*, [5]) (**a**). Example diffuse reflectance measurements of 1:1 mixture of milk and zooxanthellae (for *A. squarrosa* from mesophotic depths) in seawater for three different algal concentrations (*black*). Data were fitted using diffusion theory (*red*, see methods) (**b**).

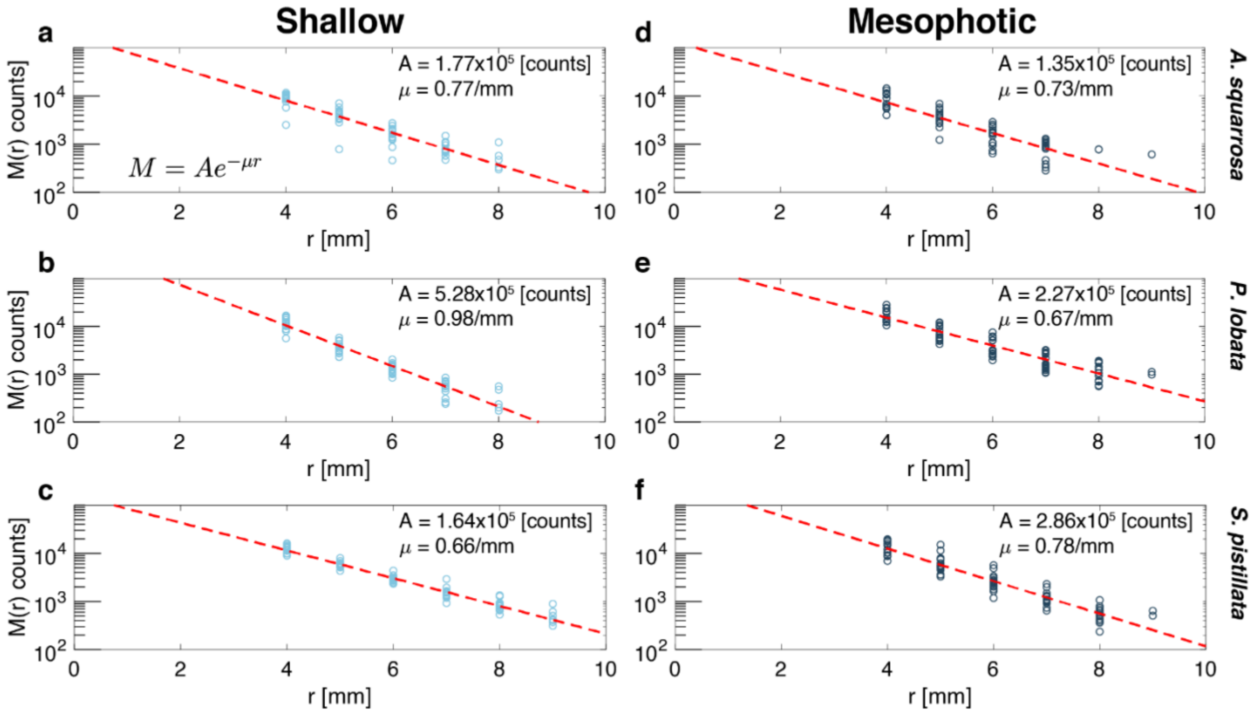


**Figure S3.** Radial escape of diffuse reflectance at 720 nm and predicted attenuation via diffusion theory. Experimental measurements on coral skeletons (hollow circles, *n* = 15-45 repeats per species per depth). Data were modeled using diffusion theory (red lines) for shallow (**a-c**) and mesophotic corals (**d-f**).

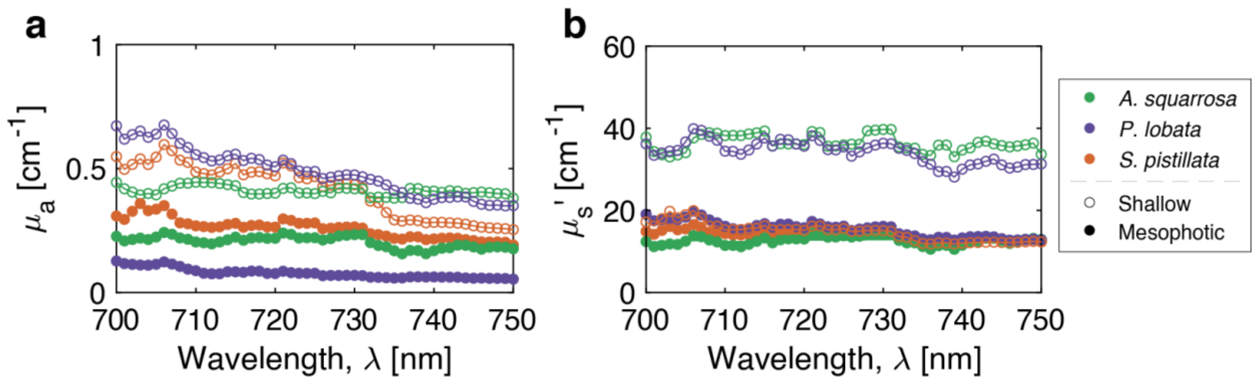


**Figure S4.** Extracted skeletal optical properties. Absorption coefficient ($\mu_a$; **a**) and scattering coefficients ($\mu_s$'; **b**) between 700-750 nm. Each color denotes a different species. Hallow and filled holes denote shallow and mesophotic depths, respectively.
